# Epigenome-Wide Study of Brain DNA Methylation Among Opioid Users and Controls

**DOI:** 10.1101/2020.11.10.377069

**Authors:** Chang Shu, David W. Sosnowski, Ran Tao, Amy Deep-Soboslay, Joel E. Kleinman, Thomas M. Hyde, Andrew E. Jaffe, Sarven Sabunciyan, Brion S. Maher

## Abstract

Opioid abuse poses significant risk to individuals in the United States and epigenetic changes are a leading potential biomarker of abuse. Current evidence, however, is mostly limited to candidate gene analysis in whole blood. To clarify the association between opioid abuse and DNA methylation, we conducted an epigenome-wide analysis (EWAS) of DNA methylation in brains of individuals who died from opioid intoxication and controls. Tissue samples were extracted from the dorsolateral prefrontal cortex of 160 deceased individuals (*M_age_* = 35.15, *SD* = 9.42 years; 62% male; 78% White). The samples included 73 individuals who died of opioid intoxication, 59 group-matched psychiatric controls, and 28 group-matched normal controls. EWAS was implemented using the Illumina Infinium MethylationEPIC BeadChip; analyses adjusted for sociodemographic characteristics, negative control and ancestry principal components, cellular composition, and surrogate variables. Epigenetic age was calculated using the Horvath and Levine clocks, and gene ontology (GO) analyses were performed. No CpG sites were epigenome-wide significant after multiple testing correction, but 13 sites reached nominal significance (p < 1.0 x 10^-5^). There was a significant association between opioid use and Levine phenotypic age (*b* = 2.24, *se* = 1.11, *p* = .045). Opioid users were approximately two years phenotypically older compared to controls. GO analyses revealed enriched pathways related to cell function and neuron differentiation, but no terms survived multiple testing correction. Results inform our understanding of the neurobiology of opioid use, and future research with larger samples across stages of opioid use will elucidate the complex genomics of opioid abuse.

## Introduction

Opioid use continues to pose significant risk to individuals throughout the globe, especially in the United States. Data from population-based epidemiological surveys from 2002-2016 in the United States revealed that approximately 30% of individuals who started heroin use met *DSM-IV* criteria for opioid dependence within one year of initiation (1). In addition, the most recent estimates from the Global Burden of Diseases, Injuries, and Risk Factors Study revealed that the global prevalence of opioid dependence was 510 people per population of 100,000, and the United States had the highest prevalence rate among all countries (1,347 persons per 100,000 population) (2). These data are alarming given the preponderance of evidence demonstrating robust associations between opioid dependence and harmful physical and psychosocial outcomes such as decreased quality of life, contact with the criminal justice system, increased risk for HIV (among injection drug users), and fatal and non-fatal overdoses (2). Given the individual and societal burden of opioid dependence, it is necessary to characterize the biological pathways to dependence. Doing so will aid in identification of specific pathways that can be targeted for treatment and intervention.

Epigenetic changes have emerged as a leading potential biological marker of drug dependence given their implications for transcription regulation and cellular reprogramming (3). Among these changes, DNA methylation (DNAm) at cytosine-guanine (CpG) sites has received the most attention in substance use research (4). Studies in animal models provide robust evidence that substance abuse causes changes in gene expression through changes in DNAm (5). Baker-Andersen et al. (6) conducted an epigenome-wide study of DNAm in the medial prefrontal cortex (mPFC) of cocaine-using mice, finding that cocaine use was associated with differential methylation in 29 regions of the mPFC and subsequent changes in gene expression. Moreover, enrichment analyses revealed that differentially methylated genes in these regions were associated with known memory and addiction pathways in the brain. Results from human studies yield similar findings (7). Hagerty et al. (8) examined epigenome-wide methylation patterns in buccal cells between alcohol use disorder (AUD) cases and controls, finding 561 hypomethylated CpG sites and 485 hypermethylated CpG sites among AUD cases compared to controls; A majority of these sites were located in genes involved in lipid metabolism, immune response, and inflammatory disease pathways. Additionally, in a second cohort of brain tissue samples, 432 of these sites (244 hypomethylated and 188 hypermethylated) were significantly correlated between buccal and brain samples, suggesting some overlap in methylation patterns across tissue type. Altogether, current evidence suggests that DNAm is a potential biomarker of substance use and has implications for broader biological functioning.

To date, epigenetic studies of opioid use have focused on candidate genes, namely *OPRM1*, which encodes for the μ-opioid receptor (9–12). This receptor plays a role in the tolerance and physical dependence of opiates (13). Candidate-epigenetic studies generally find hypermethylation of CpG sites in various regions of *OPRM1*, including promoter regions (10). Although these studies provide evidence for a potential pathway to opioid dependence in humans, they focus on a single gene, often in small sample sizes, increasing the likelihood of false positive results and limiting the detection of other genomic regions relevant to opioid use.

In the first epigenome-wide association study of DNAm of heroin users, Kozlenkov et al. (14) found three genome-wide significant differentially methylated sites in the brains of individuals who died from heroin intoxication compared to controls in tissue from the medial orbital frontal cortex (mOFC), a subregion of the mPFC that aids in goal-directed behavior and decision-making processes implicated in drug addiction (15). Recently, Montalvo-Ortiz et al. (16) conducted a genome-wide analysis of DNAm levels in whole-blood samples of opioid-dependent women and matched controls. Results revealed three significantly hypomethylated CpG sites in the *PARG, RERE*, and *CFAP77* genes, which are involved in chromatin remodeling, DNA binding, and cell survival and projection. Notably, *PARG* and *RERE* are highly expressed in multiple brain regions, and single nucleotide polymorphisms (SNPs) within *RERE* have been linked to psychiatric disorders including schizophrenia, attention deficit-hyperactivity disorder, major depressive disorder, and bipolar disorder (17). Although neither of these studies detected differential methylation in the *OPRM1* gene, they provide insight into DNAm patterns in separate tissue types (i.e., brain, blood) and elucidate the functional relevance of DNAm.

Although current evidence suggests that opioid use is associated with DNAm levels, several questions and issues remain. First, the majority of epigenetic studies focused on a single candidate gene, *OPRM1*, yet there are undoubtedly other potential genes and pathways through which opioid use contributes to dependence. Second, the majority of studies examined DNAm levels in blood or saliva, assuming that DNAm patterns in these tissues mirror levels in the brain. Although overlap in patterns of DNAm across tissues may occur, there is ample evidence suggesting that DNAm levels differ substantially by tissue type and brain region (18,19). The goal of the present study was to assess epigenome-wide DNAm levels in brain tissue of individuals who died of acute opioid intoxication and group-matched controls. The use of epigenome-wide data allows for the detection of numerous CpG sites possibly related to opioid use and dependence. Moreover, the examination of brain tissue allows us to test whether methylation patterns in the brain mirror evidence from other tissues, and to elucidate the functional relevance of DNAm in regions of the brain implicated in addiction (i.e., PFC).

## Materials and Methods

### Sample Description

Postmortem brain samples were donated to the Lieber Institute for Brain Development from the Offices of the Chief Medical Examiner of the State of Maryland (MDH protocol #12-24) and of Western Michigan University Homer Stryker School of Medicine, Department of Pathology (WIRB protocol #1126332), and one brain sample was acquired via material transfer agreement from the NIMH (donated through the Office of the Chief Medical Examiner of the District of Columbia (protocol NIMH#90-M-0142), all with the informed consent of legal next-of-kin at the time of autopsy. The present study used tissue from the dorsolateral prefrontal cortex (dlPFC) of 160 deceased individuals (*M_age_* = 35.15, *SD* = 9.42 years; 62% male; 78% White). Tables 1 and 2 provide detailed information on the study samples, which consisted of 73 individuals who died of acute opioid intoxication, 59 group-matched psychiatric controls, and 28 group-matched normal controls.

**Table 1.**
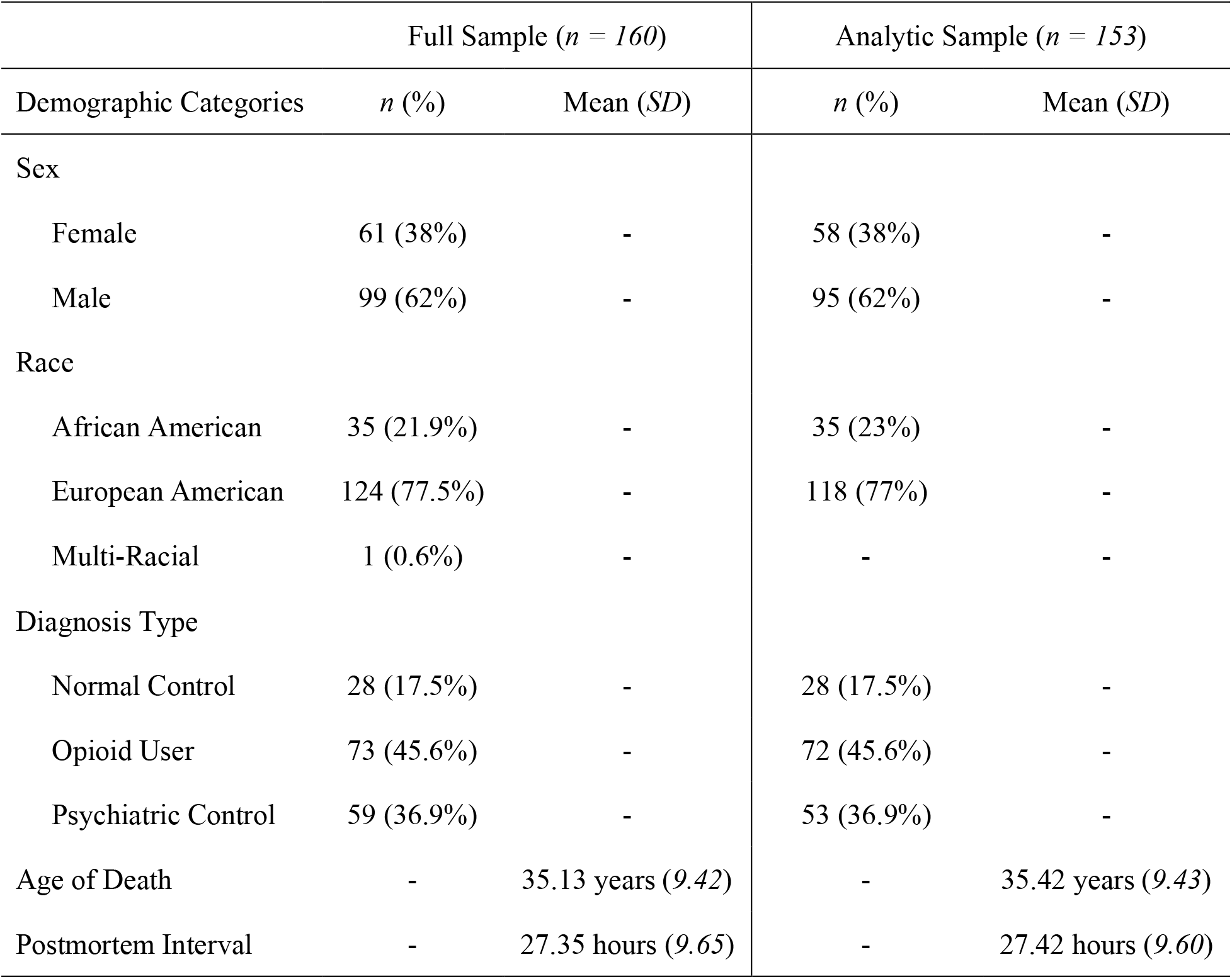
Sample Demographic Information

**Table 2.** Primary Psychiatric Diagnosis by Study Group (n = 153)

| Psychiatric Diagnosis | Psychiatric Controls | Opioid Users | Normal Controls |
| --- | --- | --- | --- |
| Attention-Deficit/Hyperactivity Disorder (ADHD) | 0 | 2 | - |
| Anxiety | 0 | 1 | - |
| Bipolar Disorder | 19 | 17 | - |
| Unspecified Bipolar Disorder | 0 | 1 | - |
| Unspecified Depressive Disorder | 1 | 6 | - |
| Major Depressive Disorder | 33 | 27 | - |
| Neurological Disorder | 0 | 1 | - |
| Obsessive Compulsive Disorder (OCD) | 0 | 2 | - |
| Eating Disorder | 0 | 1 | - |
| Posttraumatic Stress Disorder (PTSD) | 0 | 1 | - |
| Schizophrenia | 0 | 3 | - |
| Substance Use Disorder (SUD) | 0 | 10 | - |
| Total (153) | 53 | 72 | 28 |

*Note.* Two samples (one opioid user, one psychiatric control) were later removed due to mismatching predicted and observed sex, and five psychiatric controls were removed due to positive opioid tests in toxicology reports.

At the time of donation, a 36-item next-of-kin informant telephone screening was conducted to obtain medical, social, demographic, and psychiatric history. Macroscopic and microscopic neuropathological examinations were conducted on every case by a board-certified neuropathologist to exclude for neurological problems, neuritic pathology, or cerebrovascular accidents. Postmortem interval (PMI) was a calculation of hours between a donor’s time of death and the time of brain freezing. A retrospective clinical diagnostic review was conducted on every brain donor, consisting of the telephone screening, macroscopic and microscopic neuropathological examinations, autopsy and forensic investigative data, forensic toxicology data, extensive psychiatric treatment, substance abuse treatment, and/or medical record reviews, and whenever possible, family informant interviews.

All data were compiled into a comprehensive psychiatric narrative summary that was reviewed by two board-certified psychiatrists in order to arrive at lifetime DSM-5 psychiatric diagnoses (including substance use disorders/intoxication) and medical diagnoses. Non-psychiatric healthy controls were free from psychiatric and substance use diagnoses, and their toxicological data was negative for drugs of abuse. Every brain donor had forensic toxicological analysis, which typically covered ethanol and volatiles, opiates, cocaine/metabolites, amphetamines, and benzodiazepines. Some donors also received supplemental directed toxicological analysis using National Medical Services, Inc., including nicotine/cotinine testing, cannabis testing, and the expanded forensic panel in postmortem blood (or, in rare cases, in postmortem cerebellar tissue) in order to cover any substances not tested. The following substances were considered opioids: codeine, morphine, oxycodone, hydrocodone, oxymorphone, hydromorphone, methadone, fentanyl, 6-monoacetylmorphine, and tramadol.

### DNA Methylation Measurement and Preprocessing

DNA from dlFPC tissue samples was isolated with the Qiagen DNeasy kit and bisulfite converted with Zymo EZ methylation gold kit at the Johns Hopkins University Center for Inherited Disease Research. Bisulfite treated DNA was run on the Illumina Infinium MethylationEPIC BeadChip. All data cleaning and analyses were conducted in R version 3.6.1 (20). The *minfi* package was used to process raw Illumina image files into noob (normal-exponential out-of-band) preprocessed methylation beta values (21). Cell composition on percentage of positive neurons were estimated based on the method described in Houseman et al. (22) and implemented in *minfi*. Samples were tested for low intensity (*n* = 0) and inconsistency between predicted and observed sex (*n* = 2; one opioid user, one psychiatric control) and those failing were removed for quality control. Probes with low methylation intensity also were excluded (*n* = 1,208). Five additional psychiatric controls were dropped due to testing positive for an opioid in the toxicology report. The final analysis consisted of 153 samples and 864,883 probes. Principal components from negative control probes were extracted to control for technical variations (23), and surrogate variable analysis (24) was conducted to account for unknown sources of heterogeneity and remove batch effects in the data.

### Statistical Methods

#### Epigenome-wide association analysis and epigenetic age

Prior to epigenome-wide analyses, beta values were converted to M-values since the M-value distribution closer satisfies the assumption of normality in subsequent models, and the use of M-value usually leads to a better detection and true positive rate compared to the beta value (25). We used *limma* (26) to run single-site association analyses by linear regression with opioid use status (yes/no) across the epigenome for the 864,883 CpG sites, adjusting for the following covariates: diagnosis of a psychiatric disorder (yes/no), age of death, sex, postmortem interval (PMI), cell composition (i.e., % positive neurons), four negative control principal components (PCs), the top four ancestry PCs, and 13 surrogate variables detected using *sva* in R. Ancestry PCs were estimated from the 59 SNP probes profiled on the EPIC array. Next, Horvath epigenetic age and Levine phenotypic age were calculated using the *ENmix* package (27). Specifically, the difference between epigenetic/phenotypic and chronological age was obtained by subtracting the age of death of each person from their Horvath/Levine age so that positive values indicated accelerated epigenetic/phenotypic age. This value was then regressed on opioid use status and cell composition.

#### Gene ontology enrichment analysis

Gene ontology enrichment analysis was performed on probes with an FDR adjusted *p*-value < .05 using the Gene Ontology (GO) database (28) in the R package *missMethyl*, with prior correction for sampling bias (29,30). If no probes survived FDR correction, GO analyses were performed using a nominal, unadjusted significance threshold of *p*-value < .05.

## Results

### Epigenome-Wide Association Analysis

First, we conducted epigenome-wide analysis to determine if there were significant differences in methylation at individual CpG sites between opioid users and controls. Figure 1 illustrates the quantile-quantile (QQ) plot of *p*-values for the association between DNAm and opioid use. There was no evidence of inflation (lambda = 1.02). As can be seen in Figure 2, no CpG sites reached epigenome-wide significance after correction for multiple testing (red line; *p* < 5.0 x 10^-8^); however, thirteen sites reached suggestive significance using a relaxed *p*-value threshold (blue line; *p* < 1.0 x 10^-5^; see Table 3). Six of these sites were hypomethylated and the other seven sites were hypermethylated in opioid users. Among the CpG sites listed in Table 3, only cg24060527, located within netrin-1 (*NTN1*), is potentially relevant to opioid use. Evidence has linked netrin-1 activity to stimulation of the kappa opioid receptor (31,32), which is implicated in opioid dependence (33).

**Figure 1.**
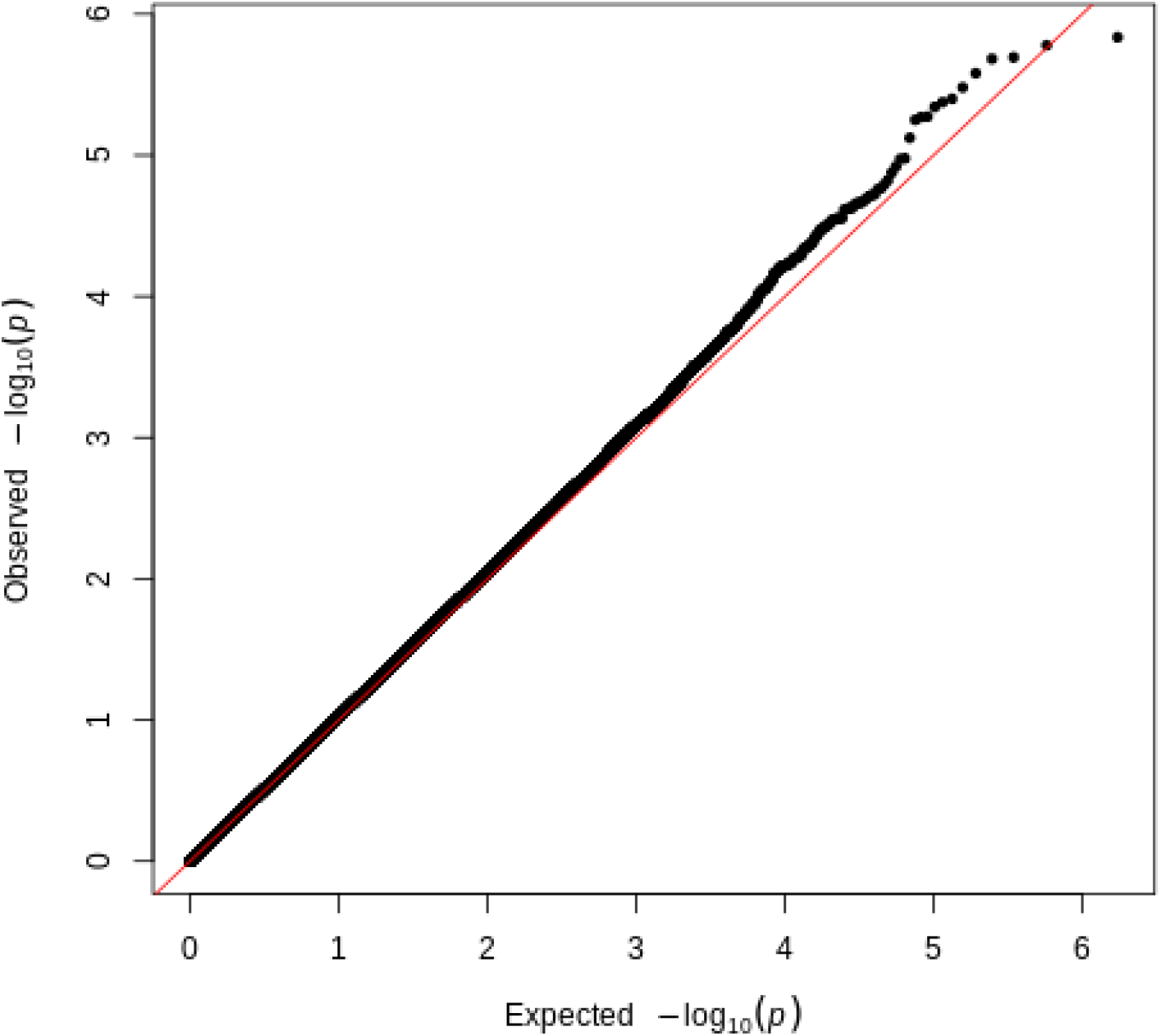
QQ Plot of p-values for the Association Between DNA Methylation and Opioid Use

**Figure 2.**
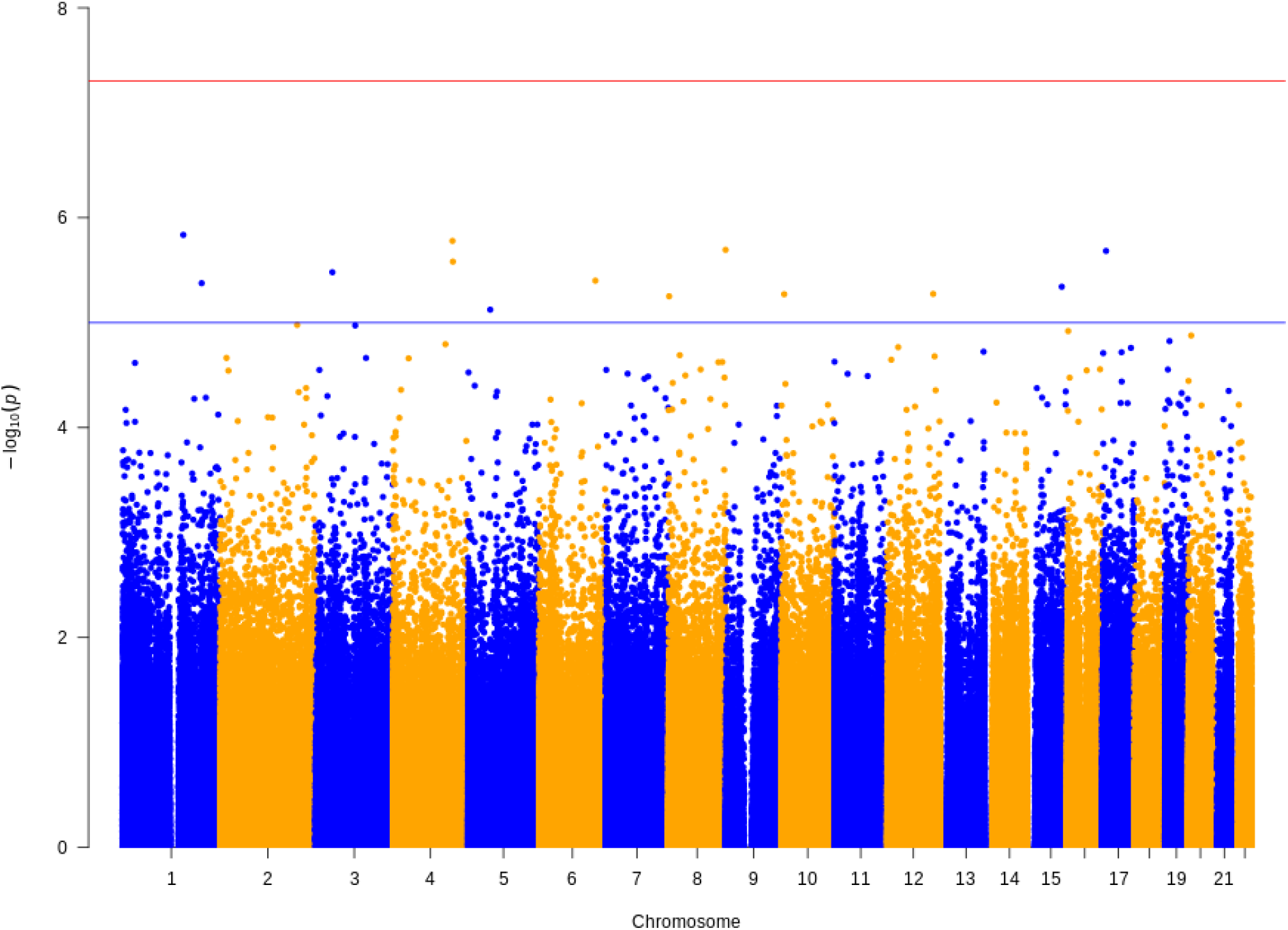
Manhattan Plot for Epigenome-Wide Association Analysis *Note*. The red line indicates genome-wide significance (*p* < 5.0 x 10^-8^) and the blue line indicates suggestive significance (*p* < 1.0 x 10^-5^). Thirteen CpG sites reached suggestive significance.

**Table 3.**
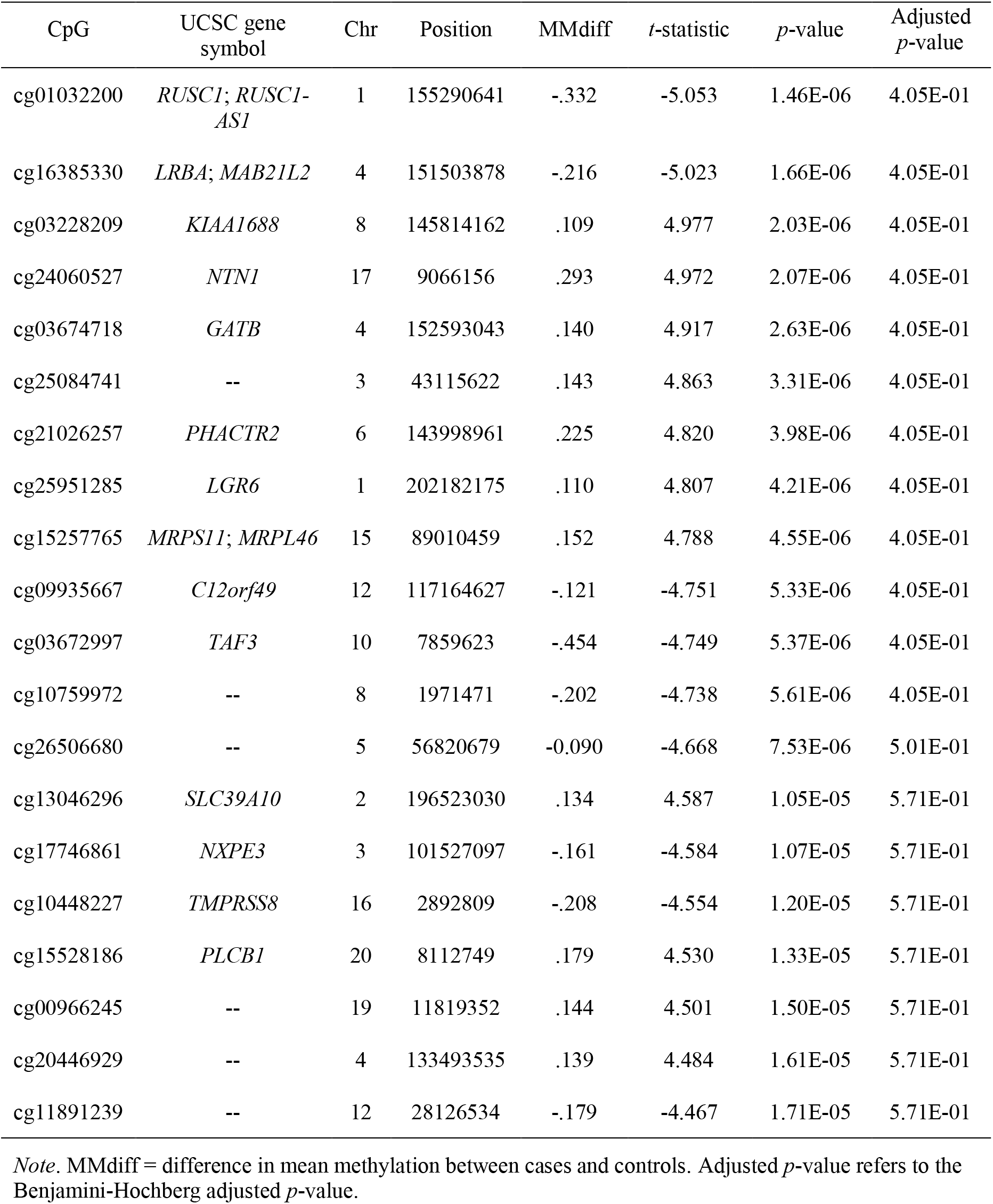
Top 20 Differentially Methylated CpG Sites

### Epigenetic and Phenotypic Age

Next, we explored the association between opioid use status and Horvath epigenetic age and Levine phenotypic age. Tests of bivariate correlations revealed that age of death was significantly associated with Horvath (*r* = .90, *p* < .001) and Levine (*r* = .69, *p* < .001) ages. Results from multivariate regressions revealed no significant association between accelerated Horvath epigenetic age and opioid use status (*b* = -.13, *se* = .67, *p* = .85), after adjusting for cell composition. There was a significant association between opioid use status and accelerated phenotypic age (*b* = 2.24, *se* = 1.11, *p* = .045), after adjusting for cell composition. Specifically, individual who died of an opioid overdose were, on average, two years phenotypically older compared to controls.

### Gene Ontology Enrichment Analyses

In order to test for case-control methylation differences at the biological system level, GO analyses were conducted for all probes with unadjusted *p*-values < .05. No terms reached significance after correcting for multiple testing (see Table 4). Examination of the top 10 GO terms did not reveal pathways relevant to opioid use.

**Table 4.**
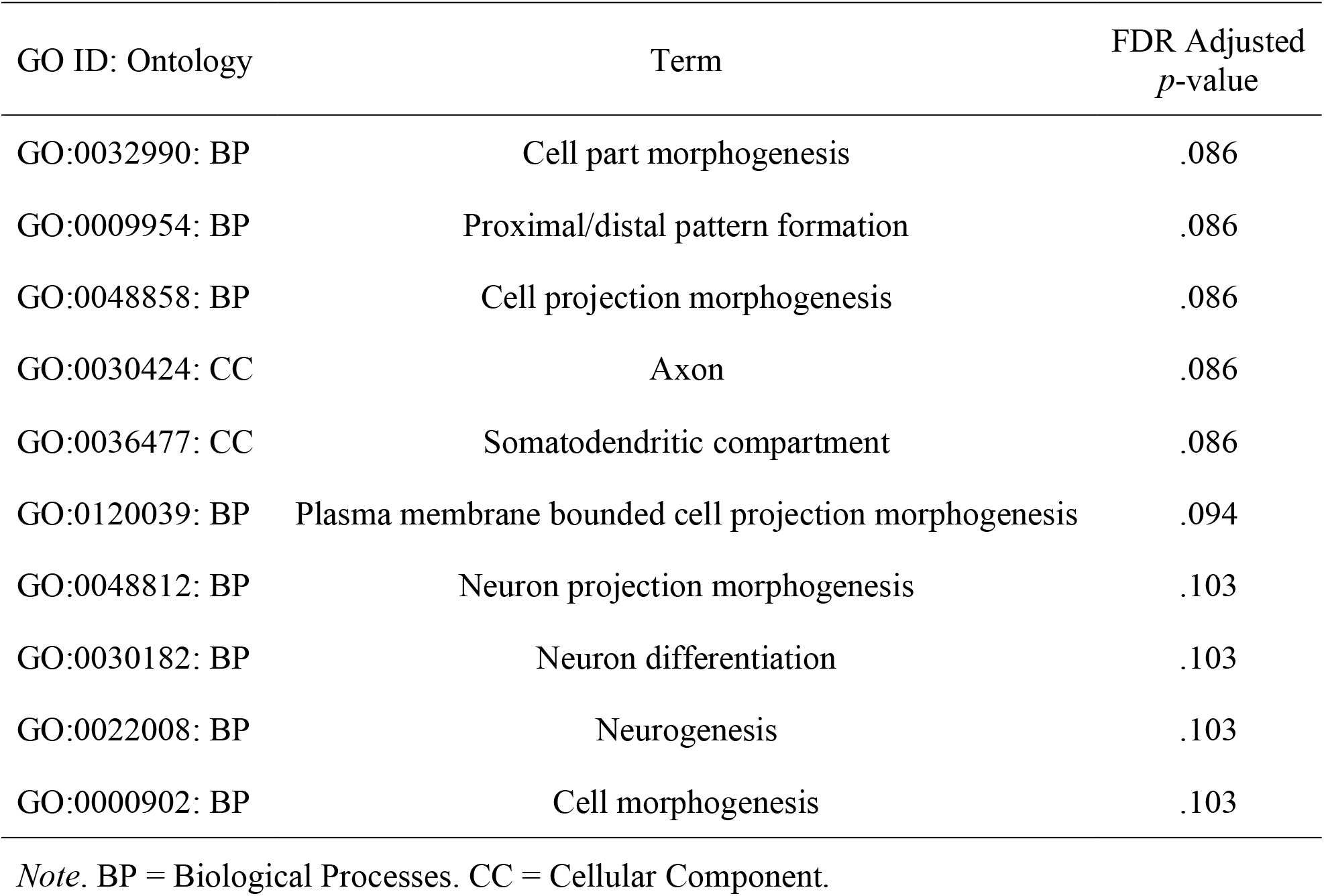
Top 10 Terms from Gene Ontology Analysis

## Discussion

The present analysis builds upon emerging literature on epigenome-wide markers of opioid abuse (16) by examining DNAm in the brains of individuals who died of acute opioid intoxication compared to group-matched controls. Specifically, the present findings inform our understanding of potential epigenetic mechanisms through which opioid use may affect pathways within brain regions involved in addiction. We observed no single-locus significant results after correction for multiple testing; however, a CpG site within *NTN-1* reached nominal significance and *NTN1* has been implicated in the stimulation of the kappa opioid receptor, which is linked to addiction and other psychiatric disorders (33). Methylation-age testing via the widely used Horvath clock did not reveal any association with opioid use status; however, analyses of Levine phenotypic age revealed older phenotypic age among those who died of an opioid overdose versus controls. Biological pathway analysis yielded multiple pathways involved in cell function and neurogenesis, although they did not survive multiple testing correction. Moreover, these were very broad pathways with limited specificity for opioid use disorder.

The significant association between opioid use and older phenotypic age supports evidence that opioid use is a risk factor for age-related negative health outcomes ranging from physical and cognitive ailments, to mortality (34). It was unexpected to not detect a significant association between opioid use status and accelerated Horvath epigenetic age, although Montalvo-Ortiz et al. (16) also failed to detect this association among a sample of women diagnosed with opioid dependence. Conversely, Kozlenkov et al. (14) found that heroin users were epigenetically younger compared to controls in their sample. Neither study assessed accelerated phenotypic age between opioid users and controls. It is possible that the divergent epigenetic age findings are due to differing samples (e.g., women only vs. men and women), examination of different tissues (whole blood vs. brain), or sampling from different brain regions, in the case of Kozlenkov et al. (14) (mOFC vs. dlPFC). Additional studies examining opioid dependence, and both accelerated epigenetic and phenotypic ages among larger samples in similar brain regions will clarify these findings.

The present study has several strengths. The primary strength of this study is the use of brain tissue in the dlPFC. This allows for testing of epigenetic markers in regions of the brain where genes of interest within addiction pathways are expressed. Second, thoughtful sampling of group-matched controls allowed us to account for potential confounders in the association between opioid use and DNAm. Third, this analysis adds to a limited literature on epigenome-wide associations between opioid use and DNAm. Although these studies have limited sample sizes and sample different tissues, each has detected CpG sites within unique genes that may have functional relevance to opioid dependence. These findings underscore the need to look beyond candidate genes to elucidate the complex etiology of opioid use and dependence.

The primary limitation of this work is the sample size. As with other investigations into the genomics of complex human traits, including substance use disorders, large sample sizes are needed to detect the effect sizes expected for the anticipated architecture of the traits. As noted by Andersen et al. (35), effect sizes for differences in DNAm between opioid users and controls are often fairly small (i.e., < 7% difference in methylation), and none of those studies used an array-based approach assessing thousands of CpG sites simultaneously, which requires larger samples to detect such small effects after correcting for multiple testing. Consequently, we and others are actively working to form consortia to analyze similar post-mortem opioid brain tissues samples together. Another limitation of this study is the lack of validation samples. Kozlenkov et al. (14) conducted the only other epigenome-wide analysis of DNAm in the brains of opioid users, examining DNAm in the mOFC. A comparison of the top 100 CpG sites from their analyses did not yield any overlapping sites. Given the dearth of studies looking at epigenome-wide associations among opioid users, it is difficult to interpret the present findings, especially given the sample size and unique population. Lastly, the cross-sectional design does not allow for direct testing of a causal relationship between opioid use and DNAm. Larger, longitudinal studies are needed to clarify the effects of opioid use on DNAm, as well as important confounders such as polysubstance use.

It is important to consider, in light of the limitations, how these results can best be used to inform the field. There are several limitations of work on substance use disorders (SUDs) in post-mortem brain tissue. First is the difficulty in distinguishing the causal role of any differentially methylated regions/pathways discovered in post-mortem tissue when compared to non-SUD control tissue since the tissue is frequently collected after years of chronic drug use. Thus, it is where in the etiologic pathways of the disorder those findings lie – causes of SUD risk or consequences of chronic opioid use disorder (OUD). Second, the utility of such findings is relatively limited with respect to clinical translation. While these findings will help elucidate our understanding of the neurobiology underlying the clinical course of OUDs, inasmuch as these findings do not overlap with findings in peripheral tissue, their potential use as a clinical biomarker is limited. Although there are limitations in the clinical utility of this work, it is important to consider how post-mortem brain SUD results can inform the field. As a clearer picture emerges from genetic studies of OUD risk and clinical course, it will be important to integrate the knowledge gained from this work.

## Conclusions

This study adds to a growing literature on genome-wide associations between opioid use and DNAm and is the first to do so using brain tissue from the DLPFC. Although no sites reached significance after correction for multiple testing, opioid users were phenotypically older compared to controls. Future work with larger samples is needed to clarify these associations.

## Funding

David W. Sosnowski is supported by the National Institute on Drug Abuse’s Drug Dependence Epidemiology Training Program (T32 DA007292-27; PI: Brion S. Maher). This research was supported by the National Institute on Drug Abuse (R01 DA039408; PI: Brion S. Maher).

## Declaration of Interest

The authors declare no competing interests.

## Acknowledgements

The authors would like to express their gratitude to our colleagues whose efforts have led to the donation of postmortem tissue to advance these studies, including at the Office of the Chief Medical Examiner of the State of Maryland, Baltimore Maryland and the Office of the Chief Medical Examiner of Kalamazoo County Michigan. We also would like to acknowledge the contributions of Amy Deep-Soboslay for her diagnostic expertise. Finally, we are indebted to the generosity of the families of the decedents, who donated the brain tissue used in these studies.

## Notes

### Competing Interest Statement

The authors have declared no competing interest.

